# Phenological responses to multiple environmental drivers under climate change: insights from a long-term observational study and a manipulative field experiment

**DOI:** 10.1101/187021

**Authors:** Susana M. Wadgymar, Jane E. Ogilvie, David W. Inouye, Arthur E. Weis, Jill T. Anderson

**Affiliations:** Department of Genetics and Odum School of Ecology, University of Georgia, Athens, GA, 30602, USA; The Rocky Mountain Biological Laboratory, Crested Butte, CO 81224, USA; Department of Biological Science, Florida State University, Tallahassee, FL, 32306, USA; Department of Biology, University of Maryland, College Park, MD, 20742, USA; Department of Ecology and Evolutionary Biology, University of Toronto, Ontario, M5S 3B2, Canada

**Keywords:** constraint, flowering onset, growing degree days, interspecific variation, photoperiod, snowmelt date, snowpack, temperature, reproductive phenology

## Abstract

Climate change has induced pronounced shifts in the reproductive phenology of plants, with the timing of first flowering advancing in most species. Indeed, population persistence may be threatened by the inability to track climate change phenologically. Nevertheless, substantial variation exists in biological responses to climate change across taxa. Here, we explore the consequences of climate change for flowering phenology by integrating data from a long-term observational study and a manipulative experiment under contemporary conditions. Dissecting the environmental factors that influence phenological change will illuminate why interspecific variation exists in responses to climate change. We examine a 43-year record of first flowering for six species in subalpine meadows of Colorado in conjunction with a 3-year snow manipulation experiment on the perennial mustard *Boechera stricta* from the same site. We analyze shifts in the onset of flowering in relation to environmental drivers known to influence phenology: the timing of snowmelt, the accumulation of growing degree days, and photoperiod. At our study site, climate change is reducing snowpack and advancing the timing of spring snowmelt. We found that variation in phenological responses to climate change depended on the sequence in which species flowered, with early-flowering species flowering faster, at a lower heat sum, and under increasingly disparate photoperiods in comparison to species that flower later in the season. Furthermore, climate change is outpacing phenological change for all species. Early snow removal treatments confirm that the timing of snowmelt governs observed trends in flowering phenology of *B. stricta* and that climate change can reduce the probability of flowering, thereby depressing fitness. Shorter-term studies would not have captured the trends that we document in our observational and experimental datasets. Accurate predictions of the biological responses to climate change require a thorough understanding of the specific environmental factors driving shifts in phenology.

## Introduction

Across ecosystems worldwide, climate change has induced shifts in the timing of critical life history transitions for a diversity of organisms, from plants to invertebrates, birds, mammals, amphibians and fish (CaraDonna *et al*., 2014, Charmantier & Gienapp, 2014, Parmesan, 2006, Parmesan & Yohe, 2003, Pilfold *et al*., 2017, Poloczanska *et al*., 2013, Wolkovich *et al*., 2012). Increased temperatures and altered precipitation dynamics have had especially noticeable effects on phenological transitions in the spring, with many species consistently now emerging, migrating, or reproducing 1-3 weeks earlier than historical averages (Amano *et al*., 2010, Bertin, 2008, Menzel *et al*., 2006, Parmesan & Yohe, 2003, Poloczanska *et al*., 2013, Sherry *et al*., 2007). In plants, shifts in the onset of flowering are among the most conspicuous and well-documented biological indicators of a changing climate (CaraDonna *et al*., 2014, Fitter & Fitter, 2002, Menzel *et al*., 2006, Parmesan & Yohe, 2003), yet we know little about which climatic factors are driving phenological responses to climate change. While flowering phenology has advanced over the past several decades for many species, substantial variation exists in responses among taxa (Mazer *et al*., 2013, Willis *et al*., 2008). Shifts in phenology alter the abiotic environment under which individuals develop and can disrupt biotic interactions when interacting species time life history events in response to different environmental conditions (Visser *et al*., 2006). Species unable to track climate change via appropriate phenological changes are at a greater risk of decline (Willis *et al*., 2008); therefore, we must explore the factors contributing to variation in responses to climate change.

Ecological and evolutionary studies typically report phenology as the calendar (ordinal) date of year, but such data may obscure the underlying biological processes that contribute to transitions between life history stages (Cook *et al*., 2012). Plants time their reproduction based on one or more proximate environment cues, including photoperiod, temperatures, the length of winter (vernalization), and moisture levels (Forrest & Miller-Rushing, 2010, Lacey, 1986). Once a cue is received, these same environmental factors influence the ensuing developmental rate. For instance, some species begin flowering only after having reached a critical photoperiod. Photoperiod correlates strongly with calendar date at a given latitude and is unaffected by climate change. Nevertheless, developmental processes initiated by photoperiod also depend on climatic factors that are changing rapidly, such as temperature and precipitation. Therefore, species that rely on fixed photoperiodic cues may now be undergoing phenological transitions under temperatures or heat sums that are now too high (Fig. 1). Species that respond solely to thermal cues may more effectively track climate change, but will experience novel photoperiods, which will influence their daily exposure to photosynthetically active radiation (Fig. 1). Climate change can decouple previously reliable seasonal conditions, which can constrain phenological responses to climate change.

**Fig. 1.**
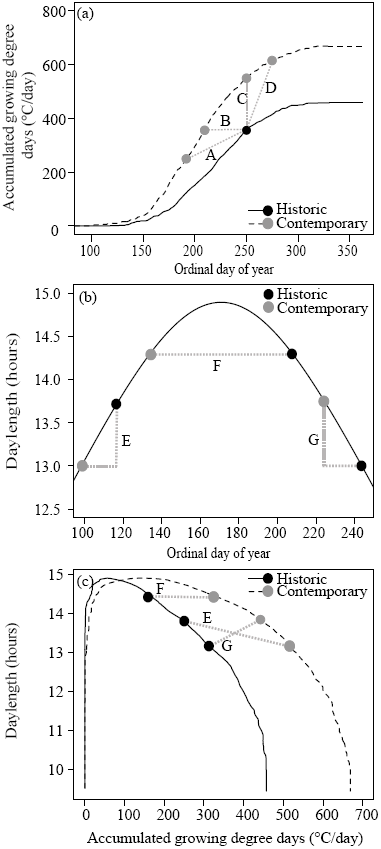
Conceptual diagrams demonstrating how climate change can decouple previously reliable seasonal cues and influence the environmental conditions that plants experience directly through shifts in climate and indirectly through shifts in phenology, (a) Accumulated growing degree days (GDD) over the growing season may shift in response to a warming climate. The historic curve shows accumulated GDD in our study region from 1975 while the contemporary curve portrays data from 2016. The reaction norms illustrate scenarios comparing the historic flowering onset dates for species (black circles) to that observed in contemporary conditions (grey circles), where we see (A) shifts towards earlier flowering onset date and decreased heat sum at flowering, (B) shifts towards earlier flowering onset date alone, (C) shifts towards increased heat sum at flowering alone, and (D) shifts towards later flowering onset and increased heat sum at flowering, (b) Shifts in the day length experienced at first flower depend on the degree of phenological change and when in the growing season a species flowers. The curve reflects the photoperiodic regime occurring in our study site. Historic shifts in flowering onset date (black circles) result in a (E) shorter, (F) equivalent, or (G) longer day length at first flower than in contemporary conditions, (c) The relationship between shifts in accumulated GDD shown in (a) and the constant photoperiods in (b). These data demonstrate how climate change and shifts in phenology may disrupt previously dependable combinations of temperature and photoperiod. The same shifts in day length depicted in (b) are reflected here and illustrate that shifts in flowering onset date for species that flower at different points throughout the growing season will result in variable combinations of accumulated GDD and day length at flowering.

Elevated temperatures during the growing season clearly influence life history traits and reproductive phenology (Wolkovich *et al*., 2012). In addition, altered winter conditions can also have profound consequences for phenological events that occur in the spring and summer. In high elevation and high latitude regions, the timing of snowmelt is a key environmental cue triggering life history transitions (Anderson *et al*., 2012). In snow-dominated regions, climate change has reduced winter snowpack and advanced snowmelt (Haye *et al*., 2013, Her *et al*., 2013, IPCC, 2014, Pederson *et al*., 2011), but the timing of frost events has not changed, exposing developing floral tissue to frost damage that would not have occurred historically under the insulation of snow (Inouye, 2008); thus damaged, plants must ‘restart’ floral development. In addition, increased evapotranspiration from spring and summer warming in concert with declining winter snowpack heightens drought conditions in the summer (Rangwala *et al*., 2012). Thus, multiple interacting environmental cues could elicit variable phenological responses among plant species that grow and flower in different portions of the growing season (Fig. 1) (Marchin *et al*., 2015).

By examining phenological trends in relation to relevant environmental drivers, contemporary studies can reveal whether climate change could outpace phenological events in natural communities. Plants could display perfect calendar day advances in flowering time that mirror climate change, such that individuals flower one day earlier for every day advance in the timing of spring. In that case, we would predict that phenological plasticity to lower photoperiod and increased frost damage would hasten biological responses to climate change. If, instead, first flowering accelerates at a rate slower than the advancement of the spring, then other environmental cues likely constrain the phenological response to climate change (Iler *et al.*, 2013). For example, an examination of first flowering data for 490 species in the south-central United Kingdom and eastern United States revealed that flowering phenology of species thought to be insensitive to climate change were in fact responding to the nullifying effects of insufficient winter chilling and warmer spring temperatures (Cook *et al*., 2012). We suggest that constraints on future phenological shifts may already be evident in current records.

We hypothesize that climate change is decoupling previously reliable environmental cues for flowering, which will constrain future phenological responses to ongoing climatic changes. Furthermore, we propose that this decoupling has more profound consequences for species that flower early in the season, when phenological shifts may increase exposure to frosts than shifts during the summer season. As a corollary, we posit that increased temperatures are not the sole mechanism underlying phenological shifts. To evaluate the environmental drivers of shifts in flowering onset dates, we first examined long-term trends in climate variables important for plant phenology. Then we analyzed shifts in the timing of first flowering for six plant species relative to plausible environmental drivers of phenological change from a long-term observational study. Finally, we conducted a manipulative experiment in contemporary environments to test the influence of snow depth and snowmelt date on flowering phenology in an ecological model plant *(Boechera stricta*, Brassicaceae). We assess whether advancing patterns of first reproduction reflect changes in the rate of floral development and the heat sum or photoperiod at which individuals flower. Alternatively, phenological shifts could simply reflect a passive response to advancing spring conditions.

The observational dataset provides an exemplary long-term record of the extent to which reproductive phenology has changed over a period of rapid climate change and can evaluate correlations between the timing of flowering and various environmental factors. However, data from natural populations cannot test causal links between environmental conditions and phenology, nor can those datasets distinguish between plasticity and possible genetic responses to environmental change. Therefore, we manipulated snowpack dynamics in our experimental study of *B. stricta* to test the extent to which climatic factors other than temperature could be implicated in changes in reproductive phenology. These experimental data enable us to evaluate whether plants can keep pace with changing seasonal dynamics via plasticity. Furthermore, this experiment can provide much needed information about the fitness consequences of ongoing climate change (Anderson, 2016) by assessing whether reduced snowpack and earlier snowmelt dates depress fitness by reducing the probability of flowering. Manipulative field experiments complement long-term observational studies by determining the specific environmental factors that promote phenological change, identifying limits to shifts in phenology, and testing whether climate change will reduce fitness. By comparing long-term records with experimental data from a common garden, we can make more robust inferences about the extent to which phenology will continue to advance with ongoing climate change.

## Methods

### Study system

We examined shifts in reproductive phenology of perennial forbs in dry rocky meadows around the Rocky Mountain Biological Laboratory (RMBL; 38’57 °N, 106’59°W; approximately 2900 m; Gothic, Colorado). Climate change has exposed subalpine meadows in this snow-dominated system to increased winter temperatures, decreased snowpack, and advancing timing of snowmelt, which has caused significant shifts in the timing of reproduction of multiple species in the plant community over the last four decades (Anderson *et al*., 2012, CaraDonna *et al*., 2014, Dunne *et al*., 2003, Gezon *et al*., 2016, IPCC, 2014, Pederson *et al*., 2011). This system is ideal for evaluating the phenological consequences of climate change owing to long-term records and experimental studies in contemporary climates. Furthermore, observations and simulations point to increased warming at higher elevations such as the Rocky Mountains (Mountain Research Initiative, 2015).

### Trends in climate

We acquired snowmelt data from two sources. First, from 1975-2016, billy barr has measured the annual date of snowmelt at the RMBL as the date of 100% bare ground in a permanent 5 x 5 m plot. Second, we estimated snowmelt dates for each plot from the long-term observational study (see below) in 2007-2016 from light intensity and temperature data recorded by HOBO pendant temperature and light data loggers (Part # UA-002-64, Onset Computer Corp, Bourne, MA, USA). We estimated snowmelt dates for the observational plots in preceding years (1973-2006) by regressing plot-level snowmelt dates on snowmelt dates measured by b. barr (*r*^2^=0.77, *F*_1,33_=110.6, *p*<l0.0001; Plot date = barr date / 0.81* 1.168 – 33.047). Previously, data on observed dates of snowmelt were combined with historic runoff values in the nearby East River to evaluate trends in the timing of snowmelt from 1935 to 2012 (Anderson et al. 2012). We update that analysis here with data from 2013-2016 (Fig. S1).

We acquired records of maximum and minimum daily temperatures from 1973-2016 from the land-based NOAA station located approximately 9 km from the RMBL (station USC00051959, https://www.ncdc.noaa.gov/data-access/land-based-station-data). Temperature data taken at the RMBL were available from permanent weather stations established in 2004 (http://www.gothicwx.org). To estimate daily temperatures in years prior to 2004, we separately regressed minimum and maximum daily temperatures taken at the RMBL against those from the NOAA station (Minimum: *r*^2^=0.88, *F*_1,4905_=36960, *p*=0, RMBL min = 0.76*NOAA min+0.46; Maximum: *r*^2^=0.85, *F*_1,4905_=29101, *p*=0, RMBL max = 0.83*NOAA max + 0.82). This 43-year record of spring temperatures lacked minimum or maximum temperatures for only 31 days. In these cases, we substituted the average of the temperatures recorded on the previous and subsequent days. We used these temperature data to estimate growing degree days for each day during the study period.

Growing degree days (GDD) are typically calculated by adjusting average temperatures by a baseline temperature below which growth cannot occur ((maximum – minimum)/2 – base). However, this method overestimates the number of growing degree days available on days when minimum temperatures are below the baseline temperature (Arnold, 1960). Here, we assign a baseline temperature of 8°C based on a resource allocation model applied to *Boechera stricta* (Colautti *et al*., 2016). In our study region, springtime minimum temperatures are consistently below this threshold (Fig. S2d). To assess GDD accurately during the spring, we modeled heat sum accumulations each day using the sine-wave method, which applies sine curves to daily maximum and minimum temperatures to approximate diurnal thermal curves (Baskerville & Emin, 1969). For both the long-term and experimental phenology data, we estimated the heat sum acquired by plants each year upon reproduction as a summation of GDD values from that year’s date of snowmelt to the ordinal date of first flower.

### Long-term phenology study

Since 1973, David Inouye and colleagues have quantified flowering phenology in the natural plant communities within 23 2 x 2 m plots near the RMBL; here, we analyze records from seven plots in dry rocky meadows ranging in elevation from 2928-2970 m over 1973-2016 (data were not taken in 1978 and 1990, *N*= 41 years). Inouye *et al.* visited each plot approximately every other day during the growing season to record the number of open flowers of each plant species (or the number of capitula with open florets for species in the Asteraceae), thereby generating an extensive long-term record of the flowering schedule of 120 species (CaraDonna *et al.*, 2014, Inouye, 2008). To examine the environmental drivers of shifts in the timing of first reproduction, we focused on six species: *Mertensia fusiformis* (Boraginaceae), *Delphinium nuttallianum* (Ranunculaceae), *Boechera stricta* (Brassicaceae), *Lathyrus leucanthus* (Fabaceae), *Vicia americana* (Fabaceae), and *Erigeron speciosus* (Asteraceae) (referred to in the results as *Mf, Dn, Bs, Ll, Va*, and *Es*, respectively). We selected these species because they are well represented in the rocky meadow plots throughout the study period, their flowering onset dates range from early spring to midsummer, and they are members of plant families representative of the communities near the RMBL. These species are pollinated by native bees, hummingbirds, or butterflies, except for *B. stricta*, which is primarily selfing.

We examined flowering phenology in relation to the timing of snowmelt by calculating the number of elapsed days from snowmelt to first flower for each species in each year, with data pooled across the seven Inouye plots. To evaluate the heat sum acquired by first flowering, we estimated the number of accumulated GDD from snowmelt to first flower. We obtained day lengths (www.sunrise-sunset.org.) for each day of the study period to examine whether the day length at first flower has changed over time.

### Experimental phenology study

To examine the mechanistic relationship between flowering time and altered winter climates, we analyzed flowering phenology data from a multiyear snow manipulation experiment. From 2014-2016, we monitored flowering phenology in experimental transplants of *B. stricta* in a common garden experiment in a dry rocky meadow near the RMBL (2891 m, 38°57.086”N, 106°59.4645”W; 2890 m elevation). *Boechera stricta* is a primarily self-pollinating perennial forb that inhabits elevations from 700-3900 m in its native Rocky Mountain range (Al-Shehbaz & Windham, 2010, Rushworth *et al*., 2011, Song *et al*., 2006). Previously, we demonstrated that *B. stricta* flowers &#x223C;13 days earlier in contemporary years than in the mid-1970s; this shift likely resulted from both plasticity and adaptation to strong directional selection (Anderson *et al*., 2012). This mustard shows plasticity in flowering phenology to temperature, winter length, and the timing of snowmelt (Anderson & Gezon, 2015, Anderson *et al*., 2012, Anderson *et al*., 2011).

In the fall of 2013 (hereafter: 2013 cohort), we planted *N*= 1691 3-month-old juvenile rosettes from 100 maternal families originating from 43 source populations into 16 randomly arrayed 2 x 1 m experimental blocks in the common garden (average = 8.4 individuals/treatment; Table S1 has exact sample sizes per genotype, source population, and numbers of individuals that flowered per growing season from 2014-2016). We replicated this experiment in the fall of 2014 (hereafter: 2014 cohort), when we planted *N=* 1839 juvenile rosettes from 154 maternal families originating from 48 source populations into 18 different experimental blocks the same common garden (average=6.0 individuals/treatment; Table S2 has sample sizes). Before transplanting these cohorts, we grew seeds from natural populations in the greenhouse for a generation to reduce maternal effects and generate maternal families consisting of selfed full siblings. The source populations spanned a broad elevational gradient (2013 cohort: 2694–3690 m; 2014 cohort: 2520–3530 m), which allowed us to investigate the phenological consequences of climate change in families with divergent evolutionary histories.

We exposed half of the transplants to contemporary climates and half to early snow removal, which reduces snowpack, accelerates the timing of snowmelt, and reduces soil moisture. In the spring of 2014-2016, we implemented the snow removal treatment, shoveling snow off of half of the experimental blocks in mid-April, when snowpack receded to 1 m deep, following our published protocol (Anderson & Gezon, 2015). To prevent damage to vegetation, we shoveled snow to 10 cm depth and allowed the remaining snow to melt naturally over the subsequent days. We placed shoveled snow outside of the vicinity of the experimental garden and shoveled a 0.5 m buffer area around each snow removal plot. We were careful to leave control plots intact. We recorded date of snowmelt (100% bare ground) for all control and snow removal plots.

As soon as snow melted, we monitored all experimental plants for flowering 2-4 times per week. At each census, we recorded the phenological status (vegetative, bolting, or flowering). We defined flowering onset as the appearance of the first flower. On reproductive plants, we measured plant height, and recorded the number of flowers, number of developing siliques (fruits), and the length of the siliques. For plants that flowered on days in between censuses, we used elongation rates of siliques and growth rates of flower production to estimate the exact ordinal date of first flowering (Table S3), as we have done previously (Anderson *et al.*, 2011, Wadgymar *et al*., 2017). For the 2013 cohort, more than half of the experimental individuals flowered during their first growing season (2014: 947 individuals), with fewer flowering in subsequent years (2015: 475 individuals; 2016: 124 individuals, Table S1). For the 2014 cohort, flowering success was lower, with 18.8% of individuals flowering in their first year (2015: 346 plants) and only 5.3% flowering in 2016 (96 individuals, Table S2).

### Data analysis

*Trends in climate.* We analyzed climatic series using both generalized additive mixed models and linear mixed models. Additive models are not constrained by the assumption of linear relationships between predictor and response variables. In this way, additive models can easily incorporate seasonal fluctuations or outlier events without compromising the fit of the predicted function. In contrast, linear models can accentuate underlying trends and yield coefficients that are easy to interpret and compare among analyses. Here, we use additive models to inspect fluctuations and anomalies in climatic or phenological data, but we focus the majority of our discussion on the linear trends uncovered from our data.

We used generalized additive models and linear models to assess changes in the date of snowmelt; the snow depth on April 1 (hereafter, snowpack); and May maximum, average, and minimum temperatures over time. We also used these models to examine the extent to which snowpack predicts the date of snowmelt, which in turn determines the spring temperatures that plants will experience. We fit additive models using unpenalized regression splines, which yield more accurate *p*-values and with the degree of smoothing selected by generalized cross-validation (Wood, 2017). For these models, significant predicted functions indicate that the data are nonlinear at some point, although they do not reveal whether there are overall increasing or decreasing trends. To gauge which portions of the predicted functions are significantly increasing or decreasing, we computed the first derivative and its confidence intervals of spline function supplied by our additive models. In all figures, portions of the dependent variable where the derivative function indicated that the slope of the predicted function was significantly different than zero are emphasized with a bold trend line. We utilize linear models to examine general trends in the data. In all linear models, we assessed the significance of quadratic terms for dependent variables, but only linear terms were retained in the final models. In both additive and linear models, we applied autoregressive autocorrelation structures and included error variance covariates to remedy residual heterogeneity where appropriate. We used the *mgcv* (Wood, 2004) package and l*m* base function in R (R Core Team, 3.2.2, *https://www.R-project.org*) to apply the generalized additive models and linear models, respectively, and used the *tsgam* package (Simpson, 2017) to estimate the derivatives of the predictive functions from the additive models.

*Phenology.* For both the observational and experimental phenology data sets, we assessed temporal trends or treatment effects on four aspects of flowering phenology: the ordinal date of first flower, the number of days from snowmelt to first flower, the number of accumulated GDD at first flower, and the day length at first flower. To meet assumptions of residual normality and homoscedasticity, we used a lognormal distribution for accumulated GDD at first flower for both observational and experimental data.

*Long-term phenology study.* We tested for temporal trends in flowering phenology from the observational study using generalized additive mixed models and linear mixed models with plot included as a random effect. We assessed whether shifts in snowmelt date and phenology were progressing at the same rate by regressing the ordinal date of first flower against the date of snowmelt for each species. To distinguish between the influence of snowmelt and thermal conditions on phenology, we include spring temperature as a covariate in these models. We used the same R packages and functions as described earlier to apply these models. We used the Benjamini-Hochberg (1995) false discovery rate procedure to correct for multiple testing and only present corrected p-values.

*Experimental phenology study.* For the manipulative experiment of *B. stricta*, we conducted repeated measures analyses in a generalized linear mixed model framework to evaluate the extent to which snow removal altered flowering phenology (Proc Glimmix, SAS ver. 9.4). We conducted models separately for each cohort. Our models included fixed effects for season, treatment, the season by treatment interaction, and repeated effects for season with an autoregressive correlation structure. We accounted for the evolutionary history of transplanted individuals through a fixed effect of source elevation and a random effect of genotype nested within population. We also included a random effect for experimental block. We focused these analyses on three response variables: flowering time (ordinal date; Poisson distribution with log link), elapsed days from snowmelt to flowering (Poisson distribution with log link), accumulated growing degree days at first flowering (lognormal distribution), and photoperiod at first flowering (normal distribution in Proc Mixed). Analyses of raw daylength data generated highly heteroscedastic residuals. Therefore, we evaluated statistical significance using randomization tests with 1000 permutations for this variable with the %rand_gen and %rand_anl macros (Cassell, 2002). Owing to different statistical distributions, we could not use a multivariate repeated measures framework to analyze all response variables separately. We applied the Benjamini-Hochberg (1995) false discovery rate to correct for multiple testing within each cohort.

Finally, for both cohorts, we conducted repeated measures logistic regression analyses to determine whether snow removal depressed the probability of flowering across growing seasons (Proc Glimmix, SAS, ver. 9.4) using the same fixed and random effects described above.

## Results

### Trends in climate

Annual measurements of snow depth on April 1 declined by 13.3 ± 0.61 cm per decade, resulting in a 33% decrease in snowpack from 1973-2016 (spline: *F*_9,30_=1.34, *p*=0.26; linear trend: *F*_1,38_=4.72,*p*=0.036, Fig. 2a, Table S4). The spring snowpack strongly predicts the date of snowmelt, with each 10 cm reduction in the snow depth on April 1 accelerating snowmelt by 2.36 ± 0.025 days (spline:*F*_9,30_=9.50, *p*<0.0001; linear trend: *F*_1,38_=89.2, *p*<0.0001, Fig. 2b, Table S4). Indeed, the ordinal date of snowmelt has advanced at a steady pace by 1.34 ± 0.053 days per decade over the past 82 years, occurring approximately 11 days earlier in 2016 than in 1935 (updated from Anderson et al. 2012; spline: *F*_9,72_=1.04, *p*=0.42 linear trend: *F*_1,80_=6.31, *p*=0.014, Fig. S1, Table S4). Although variable, spring temperatures have not changed appreciably across the study period (Fig. S2, Table S4). However, changes in the timing of snowmelt expose plants to different temporal segments of spring, with each 10-day advancement in the date of snowmelt resulting in a 0.55 ± 0.013°C decrease in the average daily temperature experienced in the first 50 days after snowmelt (spline: *F*_9,32_=2.05, *p*=0.065; linear trend: *F*_1,40_=16.8, *p*=0.0002, Fig. 2c, Table S4). Our study region is accumulating less snow over time, resulting in earlier spring seasons and a cooler thermal environment for plants.

**Fig. 2.**
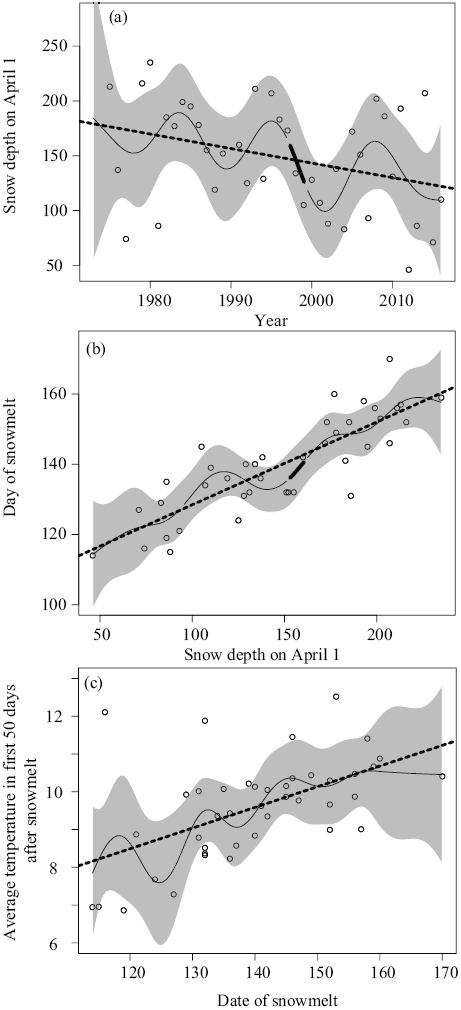
(a) Temporal trends in snow pack on April 1^st^, (b) the relationship between snowpack and date of snowmelt, and (c) the influence of the date of snowmelt on the average daily temperature experienced in the first 50 days of spring over 1973-2016 at the Rocky Mountain Biological Laboratory in Colorado. The smooth trend lines with 95% confidence intervals were derived via penalized regression splines from general additive mixed models. Portions of the predicted functions that are significantly increasing or decreasing are shown in bold (see Fig. S3). When significant, we also display the linear trend line (dashed line) from linear mixed models.

### Long-term phenology study

The ordinal dates of first flower for the six focal species were staggered throughout the growing season, with the rank order of flowering times maintained across the study period (Fig. 3a). The flowering onset dates across species fluctuate in synchrony every 8-10 years, as evidenced by the smoothing functions derived from generalized additive mixed models and their derivatives (spline: Mf *F*_10,253_=826; *Dn F*_10,261_=2217; *Bs F*_10,136_=2415; *Ll F*_10,155_=1362; *Va F*_10,40_=1717; *Es’F*_10,217_=2438;*p*<0.0001 in all cases; Fig. S3, Table S5). All species show significant advancement across the study period in flowering time as measured by ordinal day of first flowering, with greater advances in the earliest-flowering species (linear trend: *Mf F*_1,261_=56, *P*<0.0001; *Dn F*_1,269_=62, *P*<0.0001; *Bs F*_1,144_=42, *P*<0.0001; *LI F*_1,163_=18, *p*=0.0001; *Va F*_1,48_=9, *p*=0.0086; *Es F*_1,225_=21, *p*<0.0001; Fig. 3b-g, Table S5). These patterns parallel those of declining snow pack and advancing snowmelt date in this region (Fig. 2a-b). When examined as days from snowmelt to first flower, the timing of flowering has remained fairly constant in all but the earliest flowering species (Fig. 3h-n), with the onset of flowering for the early-flowering *M. fusiformis* advancing by 1.07 ± 0.043 days per decade (linear trend: *F*_1,261_=6.10154, *p*=0.024, Table S5). For all species, flowering occurred later with respect to snowmelt in a year of abnormally early snowmelt (1977, Fig. S1). There is a similar delay in flowering for the three later-flowering species in a year with an aberrantly early and cool spring (1992). The lack of a trend in flowering time when measured as elapsed days since snowmelt demonstrates that flowering generally commences at a fixed duration after snowmelt for most species.

**Fig. 3.**
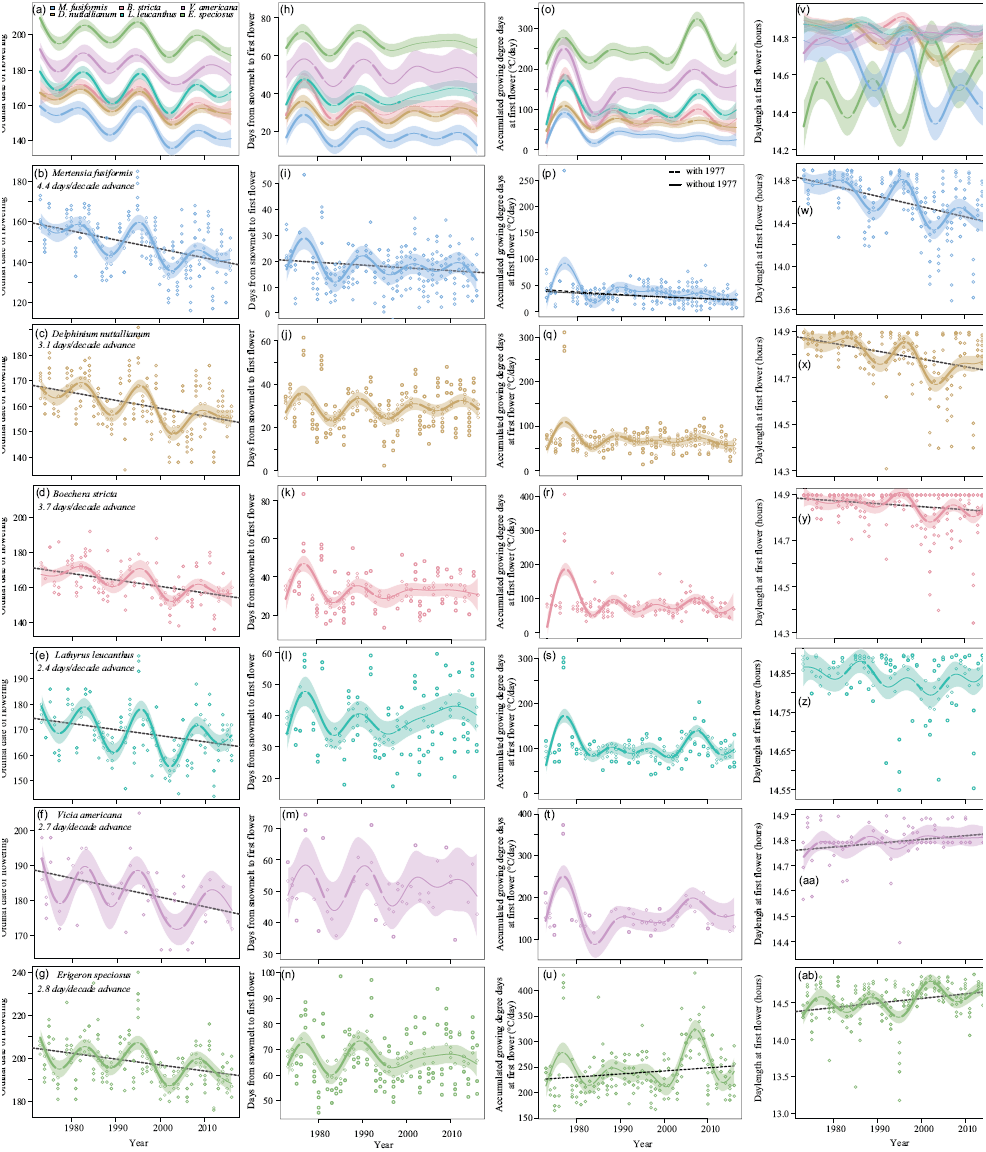
(see p. 20). Temporal trends in (a-g) ordinal flowering onset dates, (h-n) the number of days from snowmelt to first flower, (o-u) the number of accumulated growing degree days (GDD) at first flower, and (v-ab) the day length at first flower for six plant species monitored in a long-term phenology study over 1973-2016 at the Rocky Mountain Biological Laboratory in Colorado. Smooth trend lines with 95% confidence intervals were derived via penalized regression splines from general additive mixed models. The top set of panels show predicted functions for all six species together, while the raw data for each species are displayed as points in the panels beneath them in the order that species come in to flower. Portions of the predicted functions that are significantly increasing or decreasing are shown in bold (see Fig. S4). When significant, we also display the linear trend line (dashed line) from linear mixed models. Please note: Owing to the size of this figure, we have opted to provide it as the first supplemental file as well as embedding it in the manuscript file (per *Global Change Biology* guidelines). Please review the supplement for a larger version of this figure.

With snowmelt gradually occurring earlier (Fig. S1), and cooler spring temperatures in years of early snowmelt (Fig. 2c), we would expect species that are advancing their ordinal flowering onset dates to initiate reproduction at a lower heat threshold over time (Fig. 1a). However, the number of accumulated GDD at first flower have largely remained constant over time for all but the earliest flowering species (Fig. 3o-u), with peaks in 1977 (due to an exceptionally early snowmelt date, Fig. S1), and in 2007 for the later flowering species (where record maximum daily temperatures occurred on 16 of the 30 days from mid-June to mid-July). The number of accumulated GDD at flowering for the early-flowering *M. fusiformis* have decreased over the study period (spline: *F*_10,253_=9.73, *p*<0.0001; linear trend: *F*_1,261_=29.18, *p*<0.0001), and this decline is still evident when data from 1977 are excluded (spline: *F*_10,250_=12.94, *p*&10.0001; linear trend: *F*_1,258_=22.05, *p*<0.0001, Fig. 3p, Table S5).

As ordinal dates of first flower shift over time, species may encounter new photoperiodic regimes, the novelty of which will depend on their latitude and when in the growing season they flower relative to the summer solstice (Fig. 1b). The day length during the summer solstice (ordinal day 172) is 14.9 hours at our study site. Species that flower closest to the summer solstice experienced modest or no changes in day length over time despite advances in the ordinal date of first flowering (Fig. 3v-ab, Fig. S3, Table S5). In contrast, early‐ and late-flowering species are experiencing shorter or longer day lengths at flowering over time, respectively, in accordance with advances in the ordinal onset date of flowering (linear trend: *Mf F*_1,261_=45.3, p<l0.0001; *Es F*_1,225_=24.7, *p*<0.0001). The earliest flowering species, *M. fusiformis*, will continue to encounter increasingly shorter day lengths at first flower as its ordinal flowering onset date advances (Fig. 3w). In contrast, the latest flowering species, *E. speciosus*, will face more consistent day lengths as its ordinal flowering onset date shifts nearer to the summer solstice (Fig. 3ab).

The timing of snowmelt has a positive effect on ordinal dates of first flower for all species, although snowmelt is consistently advancing at a faster rate than phenology (Figure 4, Table S6). Interestingly, spring temperatures had no effect or a weak negative effect on phenology when accounting for the influence of date of snowmelt (Table S6). These results demonstrate that the date of snowmelt, and not spring temperatures, are driving shifts in phenology.

**Fig. 4.**
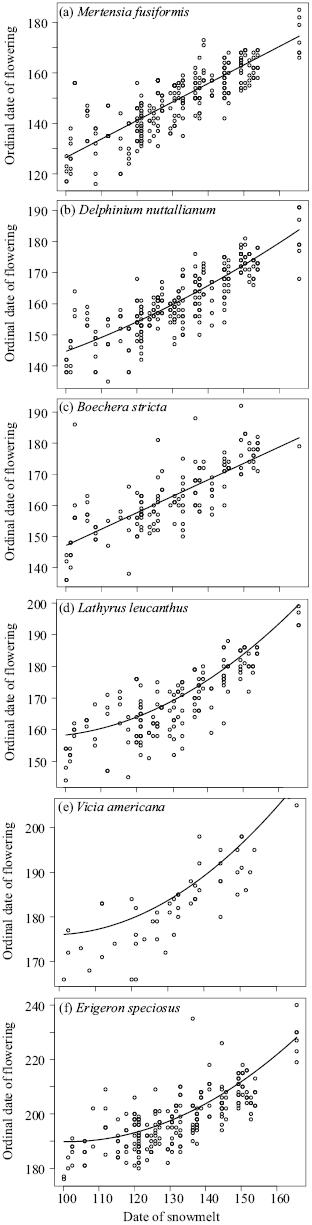
Relationships between the date of snowmelt and the ordinal date of first flower for species from the long-term phenology study in the order in which they come in to flower: (a) *Mertensia fusiformis*, (b) *Delphinium nuttallianum*, (c) *Boechera stricta*, (d) *Lathyrus leucanthus*, (e) *Vicia americana*, and (f) *Erigeron speciosus.* Linear mixed models include spring temperatures as a covariate and plot as a random effect. Linear and quadratic terms were included for both date of snowmelt and temperature.

A closer examination of phenology data for the focal species in our experimental study, *Boechera stricta*, reveals that patterns of flowering phenology are likely driven by spring snowpack. The ordinal date of first flower is positively associated with spring snowpack, with each 10 cm decrease in snowpack resulting in a 1.4-day acceleration of reproduction (spline: *F*_10,131_=2798, *p*=<10.0001;linear trend: *F*_1,139_=112.1,*p*=<0.0001; Fig. 5a, Table S4). In contrast, low snowpack delays iflowering when measured as the number of days from snowmelt to first flower, with each 10 cm decrease in snowpack producing a 1.4-day delay of reproduction relative to snowmelt (spline: *F*_10,131_=l85.98, *p*=<0.0001; linear trend: *F*_1,139_=91.15,*p*=<0.0001; Fig. 5b, Table S4). These results point to the importance of winter and early spring climate in determining the timing of flowering in this species.

**Fig. 5.**
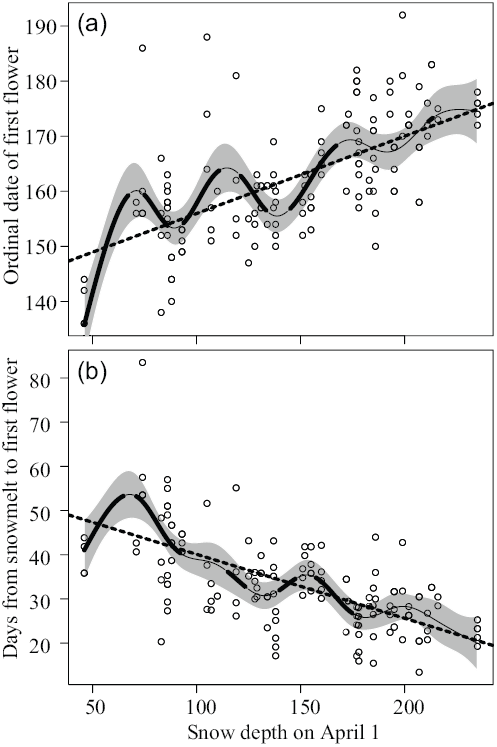
The relationship between snow depth on April 1 and (a) ordinal flowering date and (b) days from snowmelt to first flower in *Boechera stricta* over 1973-2016 in the long-term phenology study at the Rocky Mountain Biological Laboratory in Colorado. Smooth trend lines with 95% confidence intervals were derived via penalized regression splines from general additive mixed models. Portions of the predicted functions that are significantly increasing or decreasing are shown in bold (see Fig. S5). When significant, we also display the linear trend line from linear mixed models.

### Experimental phenology study

To examine the direct effects of snowpack depth and snowmelt timing on flowering phenology, we imposed an early snow removal treatment in a common garden containing populations of *Boechera stricta* from across an elevational gradient. Snow removal accelerated the timing of snowmelt to a varying extent across years, ranging from 8 days in 2014 to 30-31 days in 2015 to 13.6-18 in 2016 (Table S7). The ordinal date of first flower consistently occurred earlier in the snow removal treatment in all but the 2015 season for the 2013 cohort, with plasticity ranging from 1.7 to 7.6 days among seasons and cohorts (Fig. 6a,b; Tables S8, S9, S10). When measured as days from snowmelt to first flower, the snow removal treatment delayed flowering in 2015 (a low snowpack year) and 2016 (a moderate snowpack year), but had no effect in 2014 on the 2013 cohort (a high snowpack year; Fig. 6c,d; Tables S8, 9). Plasticity in the number of days from snowmelt to first flower in 2015 and 2016 ranged from 11.1 to 29.3 days and was greatest in 2015 for both cohorts (Fig. 6c,d; Tables S8, S9, S10).

**Fig. 6.**
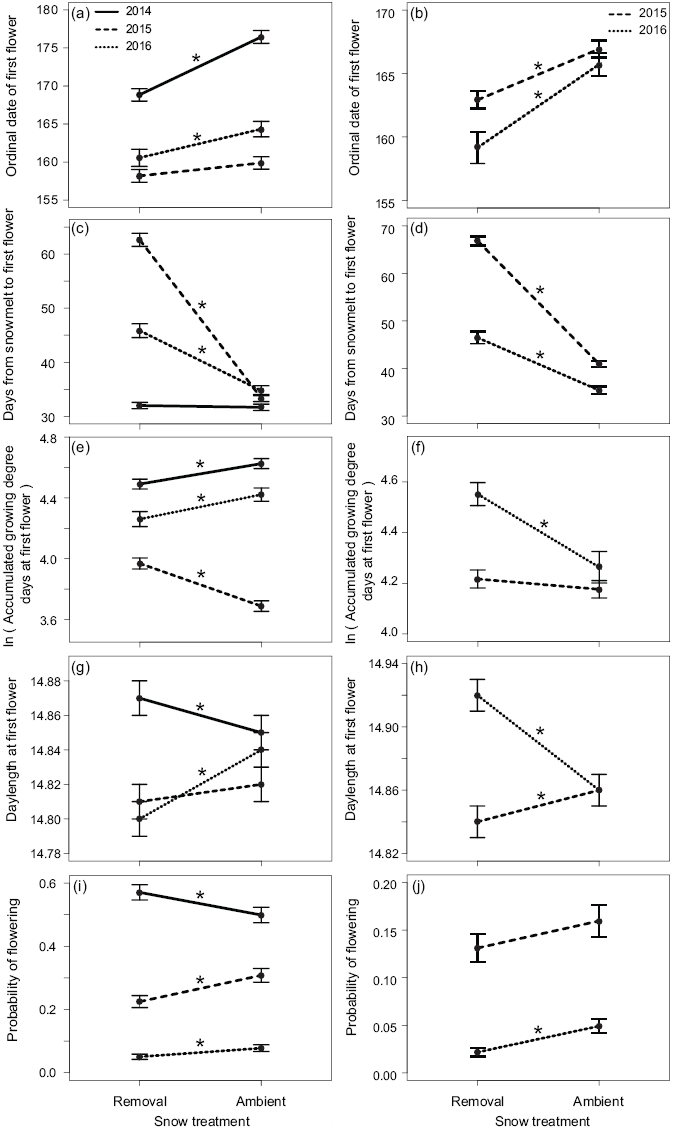
(Please see p. 25). Reaction norms depicting treatment and season effects across years from a manipulative snow-removal treatment on *Boechera stricta* in the experimental phenology study. Average (± SE) (a) ordinal date of first flower, (b) number of days from snowmelt to first flower, (c) number of accumulated growing degree days (GDD) at first flower, (d) daylength at first flower, and (e) the probability of flowering. Stars adjacent to each reaction norm are indicative of significant differences among treatments or seasons.

The effects of early snow removal on the number of accumulated GDD mirror shifts in the ordinal date of flowering onset, where years or treatments with earlier flowering correspond with a lower heat sum at first flowering (Fig. 6e,f; Table S8, S9, S10). One exception is in the 2015 growing season: For the 2013 cohort, the control treatment had significantly lower accumulated GDD than the removal; in contrast, there was no difference in heat sum across treatments in the 2014 cohort in that year. Patterns of daylength at first flowering were less consistent. In some cases, plants in the removal treatment experienced significantly lower daylengths at first flowering (2013 cohort in the 2016 growing season; 2014 cohort in 2015), whereas in other cases, we found the reverse pattern (2013 cohort in the 2014 growing season; 2014 cohort in 2016) or no difference (Fig. 6g,h; Table S8, S9, S10).

The common garden experiment also afforded us the opportunity to evaluate the probability of flowering in relation to snowmelt date. In the 2013 cohort, the probability of flowering was greater under snow removal in the first year of the study (2014), when snow removal accelerated snowmelt by only 8 days (Odds ratio 0.749, t495=0.75, p=0.0105, Table S11). For the 2014 cohort, we found no significant differences in the probability of flowering across treatments in the first growing season (2015). In contrast, both cohorts showed a significant reduction in the probability of flowering in response to early snowmelt in subsequent years (2013 cohort: 2015 Odds ratio 1.536, *t*_495_=3.45, *p*=0.0006, 2016 Odds ratio 1.585, *t*_495_=2.17, *p*=0.0307; 2014 cohort: 2016 Odds ratio 2.328, *t*_448_=3.65, *p*=0.0003; Fig. 6i,j; Table S11).

## Discussion

The ordinal dates of first flowering are advancing for species representative of the plant communities in the subalpine meadows of Colorado; this result is similar to findings from most long-term studies of flowering phenology (Amano *et al.*, 2010, Fitter & Fitter, 2002, Miller-Rushing *et al.*, 2008, Peñuelas *et al.*, 2002). Furthermore, flowering phenology is advancing at a more rapid rate in early flowering species than in those that flower later. The greater sensitivity of early-flowering species to climate change has emerged in a variety of plant taxa and across a diverse array of habitats (Dunne *et al.*, 2003, Menzel *et al.*, 2006, Miller-Rushing *et al.*, 2007, Moore & Lauenroth, 2017, Post & Nils Chr, 1999). This interspecific variation in responses to environmental change could result from pollination mode (Fitter & Fitter, 2002), increased plasticity in early-flowering species (Sherry *et al.*, 2007), variation in vernalization, and/or differences in critical photoperiod requirements between early and late flowering species (Cook *et al.*, 2012), and phylogenetic history (Mazer *et al.*, 2013). However, many potentially influential determinants of seasonal variation in phenological change remain underexplored, including variation among species in photoperiodic sensitivity and the importance of interactions among multiple aspects of climatic change (Cook *et al.*, 2012, Dunne *et al*., 2003, Matthews & Mazer, 2016).

Reporting the timing of phenological events as the ordinal dates of year on which they occur is an arbitrary, albeit convenient, choice (Sagarin, 2001). Selecting a biologically meaningful reference point from which to measure first flowering data, such as the date of snowmelt for high latitude and altitude ecosystems, could yield additional insight or entirely different conclusions about the influence of climate change on plant taxa (Bertin, 2008, Dunne *et al*., 2003). In our study, the earliest-flowering species *(M. fusiformis)* displayed a moderate advance in phenology when measured as elapsed days since snowmelt, while we detected no change for the remaining taxa. For all species, flowering commenced later with respect to snowmelt in 1977 and 1992. In these years, snow melted abnormally early, exposing plants to sub-zero temperatures. Counterintuitively, early snowmelt can result in delayed reproduction for plants that were historically insulated by snow but are now exposed to frosts in early spring (Inouye, 2008, Pardee *et al*). Results from our manipulative experiment confirm that low snowpack and early snowmelt delay flowering relative to snowmelt timing. Our data suggest that species in our system initiate flowering at a fixed time after snowmelt; therefore, shifts in the ordinal date of first flower are not the result of the development of more rapid reproductive strategies.

Variation in the date of snowmelt governs the spring temperatures that plants experience. While snowmelt is occurring earlier over time, decadal fluctuations in precipitation (Ault & St. George, 2010) and snowmelt date may prevent all but the earliest flowering species from experiencing consistent reductions in the number of accumulated growing degree days at first flower over time, although all species amass large heat sums in years with anomalous snowmelt dates. Recent extreme summer temperatures have increased the heat sum accrued at flowering for the latest species to flower *(E. speciosus)*, suggesting that climate change may differentially affect the rate at which thermal energy is accumulated for early and late flowering species. Changes in the photoperiodic regimes encountered upon reproduction also depended on the sequence in which species flowered, with early-flowering individuals experiencing increasingly disparate photoperiods and late flowering individuals experiencing increasingly stable photoperiods. Our focal species all flower in the narrow timeframe of early‐ to midsummer, when photoperiod is changing at its slowest rate. These species accumulate ~14.5 hours of daylight for photosynthesis per day, whether snowmelt occurs on day 148 (historic) or 137 (current).

Altogether, our results demonstrate that climate change may decouple historically reliable cues disproportionately among species that differ in seasonal reproductive timing. Consequently, climate change can induce flowering at inopportune times, and may even cause individuals to fail to flower. Subalpine environments have very short growing seasons, meaning that all species flower near mid-summer, when day lengths are relatively constant. Indeed, the daylength at flowering shifted by only minutes in our manipulative experiment and for the mid-summer species in the observational experiment despite significant shifts in the ordinal date of flowering onset. These changes in photoperiod are unlikely to produce biologically relevant effects on phenology for all but the earliest flowering species. In our system, continued phenological change is more likely to be restricted by unmet vernalization requirements or frost damage than by photoperiodic constraints due to the relatively constant day lengths experienced for much of the growing season. Climate change may disrupt thermal and photoperiodic regimes more in regions with longer growing seasons because greater shifts in ordinal flowering onset dates would be possible (Hülber *et al*., 2010).

The transition to flowering involves a suite of interacting developmental pathways that rely on dependable combinations or sequences of cues to signal the appropriate time to reproduce. Recent record-breaking spring temperatures in the eastern United States elicited the earliest recorded flowering onset dates for dozens of species, suggesting that phenological advances may not yet be inhibited by genetic, developmental, or physiological constraints in temperate plant species (Ellwood *et al*., 2013). Nevertheless, our observational and experimental results reveal constraints on continued phenological responses to climate change. If constraints did not exist, we would have expected plants to keep pace perfectly with climate change; that is, for every one-day advance in snowmelt, we would expect a one-day advance in first flowering. Instead, all species showed a more attenuated relationship, with flowering advancing only 0.5-0.73 days per day acceleration in snowmelt timing. This result is also evident in the manipulative study, wherein snow removal advanced snowmelt by 8-31 days, but only induced flowering to occur 1.7-7.6 days earlier than in the control treatment. Across both studies, low snowpack and early snowmelt delay flowering when measured as elapsed days since snowmelt, suggesting that plants exposed to altered snow dynamics suffer frost damage or are slower to accumulate other cues necessary for flowering. Additionally, we found that snow removal reduced flowering success in most years. These results demonstrate that climate change is outpacing plant phenological change and may be depressing fitness in our system, which could have negative consequences for future population growth rates under climate change (Anderson, 2016).

The multiyear nature of our observational and experimental studies yielded insights that would not have been apparent from shorter-term studies (Wolkovich *et al*., 2014). For all species, general additive mixed models indicated significant fluctuations in the trend of earlier flowering through time. Had we collected data for shorter periods of time (e.g., 1990-1994, or 2000-2004), we would have erroneously concluded that most species are flowering later every year, instead of the clear pattern of earlier flowering through time that we see in the longer record. Thus, temporal cycles can obscure true patterns that are not captured during a given study (Wolkovich *et al*., 2014). Multiyear field experiments are similarly valuable (Harte *et al*., 2015). Consistent with a recent meta-analysis (Anderson, 2016), we found that snow removal decreased flowering success in the experimental study, but this result only emerged after two years of snow removals had been imposed. Similarly, natural temporal variation in snowpack altered the effect of treatment on flowering phenology, resulting in some years with negligible differences in reproductive timing between control and snow removal plots.

Our results highlight that factors other than elevated summer temperatures can induce phenological shifts under climate change. In high elevation (Inouye & Wielgolaski, 2013) and high latitude (Wielgolaski & Inouye, 2013) regions, late winter and early spring conditions may play a more prominent role in structuring communities in the context of climate change, particularly for early-flowering species. By evaluating the mechanisms through which climate change alters phenology, we can make more robust predictions about biological responses to ongoing shifts in environmental conditions. Specifically, we argue that constraints will likely prevent plants from keeping pace with novel climates. As climatic conditions continue to diverge from historical values, plants in this community and others are likely to flower at times that are no longer adaptive.

## Acknowledgements

The Inouye lab thanks the many field assistants and researchers who helped collect the long-term phenology data. The Anderson lab thanks field assistants who aided in the collection of *Boechera stricta* experimental data, including Bashira Chowdhury, S. Caroline Daws, Anna Battiata, Sahana Srivatsan, Noah Workman, and Hayley Nagle. We also thank the expert skiers who completed snow removals, including Dylan Proudfoot, Turner Kilgore, Cameron Smith, Austen Beason, Ben Ammon, Alex Tiberio, Peter Innes, Kristi Haner, and Andres Esparza. We would like to thank billy barr for use of his climate data and Jennie Reithel and the Rocky Mountain Biological Laboratory for logistical support. The authors thank the National Science Foundation of the U.S. for funding these projects (DEB-9408382, IBN-9814509, DEB-0238331, DEB-0922080, DEB-1354104 to D. Inouye; DEB-1553408 to J. Anderson).

## Author contributions

D.W.I, designed and acquired funding for the longitudinal study and J.T.A. did so for the experimental manipulation. D.W.I., J.T.A., J.E.O., and S.M.W collected data. S.M.W. and J.T.A. analyzed data and wrote the first draft of the manuscript. All authors edited that draft.

## Supporting Information

### Table Captions

**Table S1.** 2013 cohort flowering success data from the experimental phenology study

**Table S2.** 2014 cohort flowering success data from the experimental phenology study

**Table S3.** Calculations of flowering onset date for the experimental phenology study

**Table S4.** Summary statistics for generalized additive (mixed) models and linear (mixed) models for analyses of trends in climate

**Table S5.** Summary statistics for generalized additive (mixed) models and linear (mixed) models for analyses of trends in phenological change from the long-term phenology study

**Table S6.** Summary statistics for generalized additive (mixed) models and linear (mixed) models for analyses of trends in phenological change from the experimental phenology study

**Table S7.** Early snow-removal treatment effects on the timing of snowmelt in the experimental phenology study

**Table S8.** Early snow-removal treatment and season effects on phenology in the experimental phenology study

**Table S9.** Least square means of phenological traits in the experimental phenology study

**Table S10.** The extent of phenotypic plasticity in phenological traits in the experimental phenology study

**Table S11.** Analyses of the probability of flowering for both planting cohorts in each treatment and across seasons in the experimental phenology study

### Figure Captions

**Fig. S1.** Trends in the date of snowmelt over time using observed dates from b. barr at the RMBL (1975-2016) and runoff measurements from the nearby East River (updated from Anderson et al. 2012).

**Fig. S2.** (a) A diagram demonstrating the contributions of daily minimum and maximum temperatures to the number of available growing degree days, and (b) Average daily maximum, (c) average, and (d) minimum temperatures in May throughout the study period.

**Fig. S3.** Derivatives and 95% confidence intervals for the predicted spline functions derived from generalized additive models for (a) temporal trends in date of snowmelt, (b) temporal and (d) the influence of the date of snowmelt on the average daily temperature experienced in the trends in snow pack on April 1^st^, (c) the relationship between snowpack and date of snowmelt, first 50 days of spring over 1973-2016 at the Rocky Mountain Biological Laboratory in Colorado (see Fig. 2, S1).

**Fig. S4.** Derivatives and 95% confidence intervals for the predicted spline functions derived from generalized additive mixed models for temporal trends in (a-f) ordinal flowering onset dates, (g-l) the number of days from snowmelt to first flower, (m-r) the number of accumulated growing degree days (GDD) at first flower, and (s-x) the day length at first flower for six plant species monitored in the long-term phenology study over 1973-2016 at the Rocky Mountain Biological Laboratory in Colorado (see Fig. 3).

**Fig. S5.** Derivatives and 95% confidence intervals for the predicted spline functions derived from generalized additive models for the relationship between snow depth on April 1 and (a) ordinal flowering date and (b) days from snowmelt to first flower in *Boechera stricta* over 1973-2016 in the long-term phenology study at the Rocky Mountain Biological Laboratory in Colorado (see Fig. 5).

